# Higher prevalence of sacbrood virus in highbush blueberry pollination units

**DOI:** 10.1101/2024.03.20.585971

**Authors:** Alison McAfee, Sarah K. French, Nadejda Tsvetkov, Heather Higo, Julia Common, Stephen F. Pernal, Pierre Giovenazzo, Shelley E. Hoover, Ernesto Guzman-Novoa, Robert W Currie, Patricia Wolf Veiga, Ida M. Conflitti, Mateus Pepinelli, Lan Tran, Amro Zayed, M. Marta Guarna, Leonard J. Foster

**Author notes:** Authors contributed equally (M. Marta Guarna, Leonard J. Foster).

## Abstract

Highbush blueberry pollination depends on managed honey bees (*Apis mellifera*) for adequate fruit set; however, beekeepers have raised concerns about poor health of colonies after pollinating this crop. Postulated causes include agrochemical exposure, nutritional deficits, and interactions with parasites and pathogens, particularly *Melisococcus plutonius*(the causal agent of European foulbrood disease), but other pathogens could be involved. To broadly investigate common honey bee pathogens in relation to blueberry pollination, we sampled adult honey bees from colonies at time points corresponding to before (t1), during (t2), at the end (t3), and after (t4) highbush blueberry pollination in British Columbia (BC), Canada, across two years (2020 and 2021). Nine viruses as well as *M. plutonius*, *Vairimorpha ceranae* and *V. apis* (formerly *Nosema ceranae* and *N. apis*) were detected by PCR and microscopy and compared among colonies located near and far from blueberry fields. We found a significant interactive effect of time and blueberry proximity on the multivariate pathogen community, mainly due to differences at t4 (corresponding to roughly six weeks after the beginning of the pollination period). Post-hoc comparisons of pathogens in near and far groups at t4 showed that detections of sacbrood virus (SBV), which was significantly higher in the exposed group, was the primary driver. The association of SBV with highbush blueberry pollination may be contributing to the health decline that beekeepers observe after pollinating this crop, likely in combination with other factors.

## Introduction

Highbush blueberry (*Vaccinium corymbosum*) is a major crop in Canada and the United States (Protzman 2021, Agriculture and Agri-Food Canada 2023) and fruit production is enhanced by insect pollination, which increases fruit weight by approximately 75% (Eeraerts et al. 2023). Many native bee species, especially those capable of “buzz-pollination” (vibrational disturbance of pollen) are more efficient pollinators of blueberry flowers than honey bees (*Apis mellifera*), per individual (Rogers et al. 2013, 2014, Hoffman et al. 2018, Cortés-Rivas et al. 2023). However, honey bees – though not native to the Americas – can be stocked at high densities and moved to specific locations, making them a common commercial pollinator (Isaacs and Kirk 2010, Gibbs et al. 2016, Hoffman et al. 2018). British Columbia accounts for 95% of Canada’s total highbush blueberry production, with particularly high densities of blueberry fields occurring in the Fraser Valley (Agriculture and Agri-Food Canada 2023).

Blueberry pollination contracts are an important source of income for beekeepers (Bixby et al. 2023). However, some beekeepers and researchers have raised concerns that colony health deteriorates after engaging in highbush blueberry pollination (Wardell 1982, Higo et al. 2019, Thebeau et al. 2022, Thebeau et al. 2023). Several sources have postulated that European foulbrood (EFB, caused by *Melissococcus plutonius*), exposure to fungicides, nutritional deficits, and their interactions could be the underlying causes of poor colony health outcomes perceived to be associated with highbush blueberry pollination (Wardell 1982, Graham et al. 2021, Graham et al. 2022, Thebeau et al. 2023).

Fungicides and their adjuvants (non-active ingredients that enhance pesticide performance, such as surfactants) can increase honey bee susceptibility to pathogens, such as *Vairimorpha ceranae* (a microsporidian parasite, formerly known as *Nosema ceranae* (Tokarev et al. 2020)) (Pettis et al. 2013), viruses (Degrandi-Hoffman et al. 2015, Fine et al. 2017, O’Neal et al. 2019), and, in some cases, *M. plutonius* (Thebeau et al. 2023). Although honey bees can also be exposed to fungicides and other agrochemicals while foraging on non-blueberry plants (Pettis et al. 2013, Graham et al. 2022), these interactions between fungicides and pathogens broadly highlight the utility of analyzing multivariate pathogen responses in crop exposure trials.

EFB is a bacterial disease affecting honey bee larvae (Forsgren 2010, Lewkowski and Erler 2019). It is thought to be a highly prevalent, but opportunistic pathogen with symptoms tending to appear during spring in association with additional stressors (Wardell 1982, Bailey and Ball 1991, Fünfhaus et al. 2018, Grant et al. 2021). EFB presentation is sometimes associated with blueberry pollination (Gregoris A 2019, Grant et al. 2021), but a recent study in Michigan calls that association into question (Fowler et al. 2023). In addition, while deformed wing virus loads appear not to influence a colony’s likelihood of developing EFB (Fowler et al. 2023), relationships with other diseases have not been fully explored.

Here, we conducted an experiment evaluating twelve pathogens, which are among those most commonly observed in honey bee colonies (Evans and Schwarz 2011, Borba et al. 2022), in colonies at different proximities to highbush blueberry fields in British Columbia. Colonies were placed in or near highbush blueberry fields (“near” sites) or at least 500 m away from blueberry fields (“far” sites) and sampled over time, starting immediately before the highbush blueberry pollination period and ending approximately two weeks after the pollination period. We hypothesized that pathogen profiles in colonies placed near and far from blueberries would differ over time, and that *M. plutonius* detections in particular would be higher in colonies placed near blueberries after pollination concluded.

## Methods

### Honey bee colonies and blueberry field sites

The honey bees used in the highbush blueberry study consisted of 40 colonies in 2020 and 40 different colonies in 2021. In 2020, the experimental colonies were produced on-site from overwintered colonies headed by queens overwintered in British Columbia, Canada. The donor colonies were treated for *Varroa* mites in early April with Formic Pro™ in according to manufacturer’s instructions. In 2021, experimental colonies were produced from nucleus colonies headed by locally mated, overwintered BC queens and allowed to grow until the beginning of the experiment. Each colony was initially housed in a single-box, deep Langstroth hive, with additional boxes added as needed to suppress swarming as the population expanded over time. Colony sizes followed conventions used for blueberry pollination units (minimum of 4 frames of brood and 8 frames of adult bees), and colonies received no supplemental feeding or medications during the course of the experiment.

Sites near and far from blueberry fields were chosen such that no two sites were within 3 km of one another, with the exception of two exposed sites which were 2 km apart. Before the pollination period, the colonies destined for different exposure sites were held in the same four yards in 2020 or three yards in 2021; then, colonies at each initial site were randomly assigned to five sites “near” highbush blueberry or five sites “far” from highbush blueberry with equal numbers of colonies (n=4) at each site (where each site formed one replicate). The colonies were moved to near and far sites on the same day where possible, or within a maximum of 24 hours of each other, when the blueberry bloom reached approximately 5-10% (as is conventional for pollinating this crop). See **Table 1** for a timeline of sampling and moving events. In both years, colonies were then relocated to post-pollination yards. In cases where some colonies remained at the same location for consecutive sampling events but other colonies were moved, a “sham” movement procedure was followed, such that all colonies were treated as if they were moving (*i.e*., they were screened, strapped, and driven in the back of a vehicle for similar durations of time), except that the colonies remaining in place were dropped off at the same location they were at previously. See **Figure 1** for a map of field sites at each time point.

**Figure 1.**
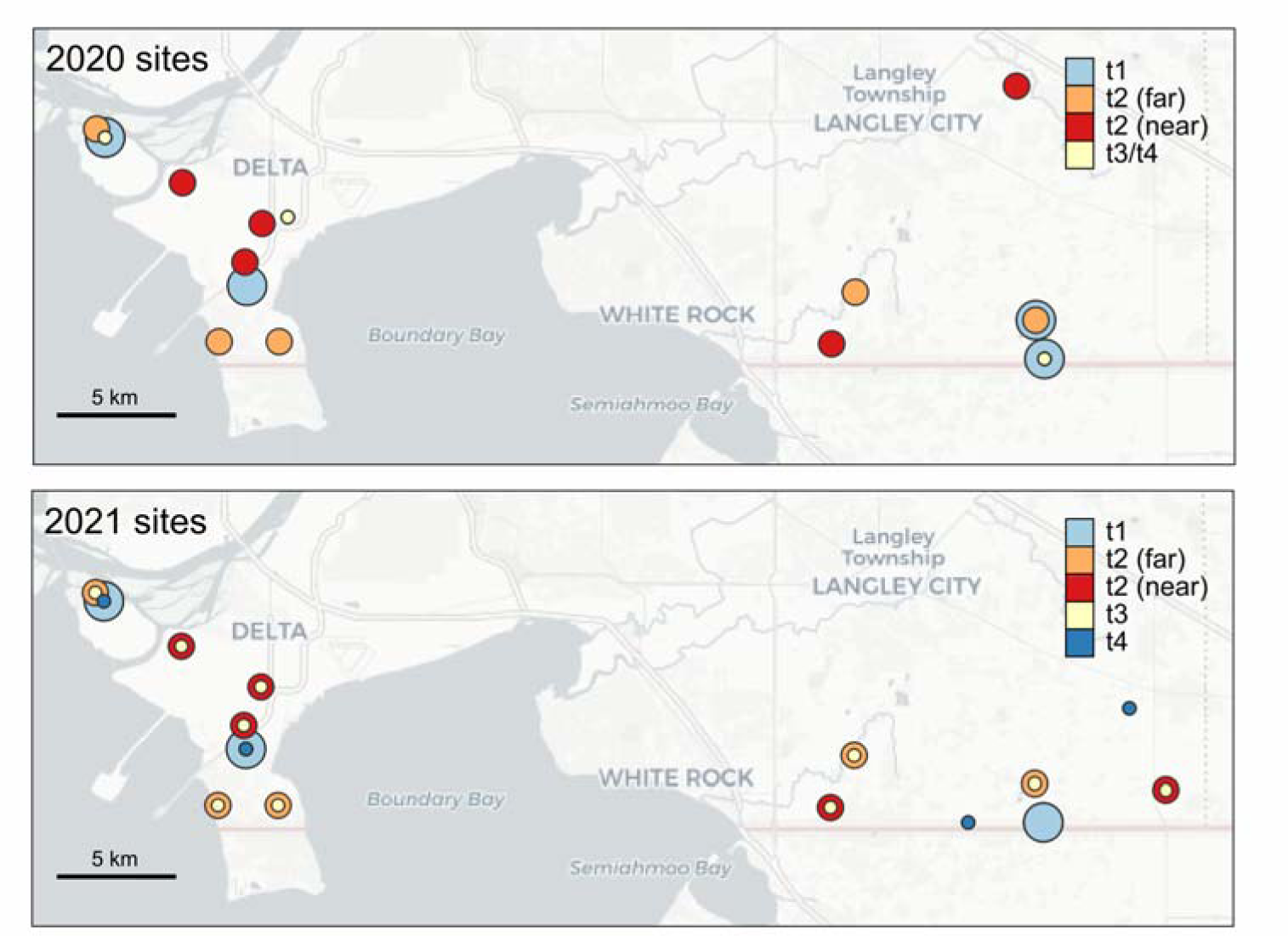
Experimental sites. N = 10 pooled samples (5 sites near highbush blueberries and 5 sites far from highbush blueberries), representing 40 colonies, were evaluated in 2020. T1, t2, t3, and t4 samples were taken at approximately two-week intervals. Large blue dots represent pre-pollination holding yards where colonies were maintained prior to prior to being moved into blueberry fields at t1. Orange and red dots show the locations of near and far site locations. Small yellow dots show the locations of colonies at t3 sampling, after the pollination period, and small dark blue dots show locations at t4 sampling. In 2020, t3 samples were taken immediately after moving out of pollination yards.

**Table 1.** Sampling dates for the highbush blueberry field experiment.

| Year | Activity | Sampling date | Days from latest t1 |
| --- | --- | --- | --- |
| 2020 | t1 sampling | April 27-28 | 0 |
|  | Move to blueberry exposed/control yards | April 29 | 1 |
|  | t2 sampling | May 11 | 14 |
|  | Move to post-pollination yards | May 25 | 27 |
|  | t3 sampling | May 26 | 28 |
|  | t4 sampling | June 11 | 44 |
| 2021 | t1 sampling | April 23 | 0 |
|  | Move to blueberry exposed/control yards | April 24-25 | 1-2 |
|  | t2 sampling | May 10 | 17 |
|  | t3 sampling | May 21 | 28 |
|  | Move to post-pollination yards | May 22-23 | 29-30 |
|  | t4 sampling | June 2 | 40 |

### Landscape data for t2 sites

We obtained land cover data for British Columbia from Agriculture and Agri-Food Canada’s (AAFC) 2020 and 2021 Annual Crop Inventory (Agriculture and Agri-Food Canada 2020, 2021). We used QGIS (version 3.26.0) to identify land cover categories and area within a 1500-m buffer surrounding each site. These categories, as defined by AAFC, included: barley, beans, blueberry, broadleaf, coniferous, corn, cranberry, exposed land or barren (excluding fallow agriculture), fallow, grassland, greenhouses, hops, mixedwood, nursery, oats, orchards, other berries, other crops, other fruits, other vegetables, pasture or forage (may include tame grasses, alfalfa, and clover), peas, potatoes, shrubland, sod, urban or developed (predominantly built-up/developed land, including associated vegetation), vineyards, water (freshwater and/or marine), and wetlands. These data were then sorted into categories comprising ≥1% and <1% of the total 1500 m radius area around each site, and areas for those with <1% coverage were summed and labelled “other”. These data are available in **Supplementary Data 1**.

### Sampling methods

Each colony was sampled at four time points: before pollination (t1), during pollination (t2), at the end of pollination (t3), and two weeks after the end of pollination (t4). Each time point was approximately two weeks apart (see **Table 1** for exact dates). Samples at t1 were taken immediately before moving colonies to their respective near and far field sites, t2 samples were taken at peak bloom, t3 samples were taken at the end of pollination (immediately before moving to t4 sites in 2021 and immediately after moving to t4 sites in 2020), and t4 samples were taken two weeks after the end of the pollination period. At each time point, adult bees were sampled from an open brood frame into a 50 mL conical tube, placed immediately on dry ice, and transported to the University of British Columbia laboratory where they were stored at -70°C until all samples were collected. Bees were also sampled into a urine vial (∼120 mL, with an average of 260 bees per sample), which was then filled with 70% ethanol for conducting mite counts by the alcohol wash method as previously described (Borba et al. 2022). Samples for pathogen analysis were shipped on dry ice to York University (Toronto, ON, Canada), where they were centrally stored at -70°C and then shipped to National Bee Diagnostic Center (NBDC) in Beaverlodge, AB, Canada. Before submission of the samples to the NBDC for pathogen analysis, bees from colonies belonging to the same site and replicate were pooled to create two composite samples (15 bees per colony, 60 bees in total in each sample). The final replication was therefore n = 5 near and n = 5 far replicates in each year, where each replicate is represented by a pool of bees from four colonies at each replicate site.

### Pathogen detection

From the pooled samples described above, the NBDC conducted pathogen testing of Israeli acute paralysis virus (IAPV), deformed wing virus A (DWV-A), varroa destructor virus (VDV; also known as deformed wing virus B or DWV-B), acute bee paralysis virus (ABPV), Kashmir bee virus (KBV), chronic bee paralysis virus (CBPV), black queen cell virus (BQCV), sacbrood virus (SBV), *M. plutonius, V. apis* and *V. ceranae* (Tokarev et al. 2020)). For *Vairimorpha* spore counts, 60 bees were homogenized in 60 mL of 70% ethanol and a 6 µL aliquot was loaded into a counting chamber (hemocytometer) with Thoma ruling and a cell depth of 20 µm. Samples were examined for *Vairimorpha* spores twice under phase contrast, at 400 x magnification (Eclipse Ci-L, Nikon, Tokyo, Japan), with spores counted from all 16 squares of the gridded area. The average number of spores per bee are reported (**Supplementary Data 2**).

For *V. ceranae*, *V. apis,* and *M. plutonius* detection, DNA was extracted from a 200 µL aliquot of the homogenized sample described above. The samples were pelleted by centrifugation, the liquid was aspirated, and the pellet was allowed to dry at room temperature. Genomic (g)DNA was extracted and purified using the NucleoSpin®Tissue kit following the manufacturer’s instructions (Macherey-Nagel Gmbh & Co. KG, Düren, Germany). Primers used for the detection of all pathogens are found in **Supplementary Data 3.**

Identification of *Vairimorpha* species was performed by end-point PCR using AccuStart™ II PCR Supermix (Quanta Bioscience, USA). Amplification assays were performed using 60 ng of gDNA in an Applied Bioscience Veriti 96-well Thermal Cycler (Applied Bioscience, Singapore). PCR conditions were 5 min at 95°C for initial denaturation/enzyme activation followed by 35 cycles of 1 min at 94°C, 1 min at 58°C and 1 min at 72°C, with a final extension of 10 min at 72°C. Amplicons were visualized by gel electrophoresis.

*M. plutonius* was detected by performing quantitative PCR using SsoAdvanced™ Universal Probe (Bio-Rad Laboratories, Hercules, USA). Amplification assays were performed in triplicate with 60 ng of gDNA on an ABS 7500 Fast System (Applied Biosystems, Foster City, USA) using a program of 2 min at 50°C, 10 min at 95°C, and 40 cycles of 15 s at 95°C and 1 min at 60°C. B-actin was used as the reference gene. Standard curves were prepared from plasmids harboring the target amplicons with copy numbers diluted serially over five orders of magnitude. Results were analyzed with the 7500 Software v2.3 and exported to calculate copy numbers per bee.

All viruses were detected and quantified using RT-qPCR following sample preparation methods as previously described (Borba et al. 2022). Briefly, total RNA was extracted from the second pooled sample of 60 bees using a NucleoSpin®RNA kit (Macherey-Nagel Gmbh & Co.), cDNA was synthesized from 800 ng total RNA using the iScript cDNA synthesis kit (Bio-Rad Laboratories), and reactions were assembled with sSoAdvanced™ Universal SYBR® Green Supermix (Bio-Rad Laboratories). Each virus was tested in triplicate for each sample using 3.75% of the cDNA reaction product, and absolute quantitation was performed by comparing sample values against a standard curve (generated from serially-diluted, amplicon-containing plasmids). PCR conditions were 3 min at 95°C for initial denaturation/enzyme activation followed by 40 cycles of 10 seconds at 95°C and 30 seconds at 60°C (except IAPV, where annealing/extension was 45 seconds at 60°C). Specificity was checked by performing a melt-curve analysis (65-95°C with increments of 0.5°C and 2 s/step). Results were analyzed with the CFX Manager™ Software and exported.

### Multivariate community analysis

Statistical analyses were conducted in R (version 4.3.0) using R Studio (version 2023.09.1+494) (R Core Team 2023). To determine if exposure to blueberry fields affected the composite pathogen profiles over time, we combined 2020 and 2021 data and conducted a PERMANOVA analysis using the adonis2 function within the R package vegan (version 2.6-4) (Oksanen et al. 2022). Similarity was calculated using the Jaccard index, since some of the pathogens (*V. ceranae*, *V. apis,* and *M. plutonius*) are limited to a presence/absence data type. We first evaluated the pathogens at time point 1 to determine if the multivariate pathogen structure among colonies destined to be distributed to different site types was indeed similar. This model included a pathogen matrix of twelve response variables (nine viruses, *M. plutonius*, *V. ceranae*, and *V. apis*), and explanatory variables of site type (levels: near and far), and year (levels: 2020 and 2021). Varroa load was initially considered in the model, but was dropped as it was not a significant explanatory factor. The pathogen data and sample metadata are available in **Supplementary Data 2**.

After confirming that site type groups were indeed similar before being moved into their respective control and exposed locations, we tested for effects of exposure on multivariate pathogen structure for the remaining time points. This model included the same twelve response variables, and explanatory factors site type (levels: near and far), and time (levels: t2, t3, and t4), as well as their interaction. We conducted a restricted permutation test, which only allows samples to be permuted within replicates, to account for repeated measures over time. Varroa load was again initially considered, but dropped from the final model as it was not a significant explanatory factor. Because an interactive effect between site type and time point was identified, we conducted *post hoc* tests at each time point individually to identify the patterns driving the interaction, again using a reductive modelling approach similar to the examples above.

### Visualization of data and assessment of specific pathogens

To visualize the multivariate data at each time point, we used the metaMDS function (package: vegan, version 2.6-4; specifying k = 2, trymax = 500, and distance = “jaccard”) (Oksanen et al. 2022) and plotted the resulting scores using ggplot2 (Wickham 2016). Differences in individual pathogen detections at t4 were also evaluated statistically using a generalized linear model (package: stats, base R) (R Core Team 2023) with a binomial vector of pathogen presence/absence as the response variable, site type and year as fixed factors, and a binomial distribution specified. Here and subsequently, we ensured appropriateness of fit by checking simulated residual plots, as enabled by the package DHARMa (version 0.4.6) (Hartig 2022). *M. plutonius* and SBV detections were also analyzed over time using a generalized linear mixed effects model (package: lme4, version 1.1-33) (Bates et al. 2015), with the respective binomial presence/absence vector as the response variable, year as a fixed factor, site type and time point as an interactive term, and replicate as a random intercept term. In both cases, year was not influential and was dropped from the final model. Summary statistics for main effects were extracted using the Anova function (package: car, version 3.1-2) (Fox and Weisberg 2019).

## Results

### Landscape descriptions

Although our original goal was to compare colonies near and far from blueberries, with far colonies being at least 1.5 km away from blueberry fields, due to the extreme prevalence of blueberries where this experiment took place (the Fraser Valley of British Columbia), choices for sites far from blueberries were limited. Eight out of ten far sites still had some blueberry cropland occurring within 1.5 km of the site; however, only 3 out of 10 had blueberry coverage representing > 1% of the total 1.5 km radius area (**Figure 2**). While the far sites were not completely devoid of blueberry cropland, they are still biased toward lower coverage and the blueberry fields that did exist were, by definition, farther away.

**Figure 2.**
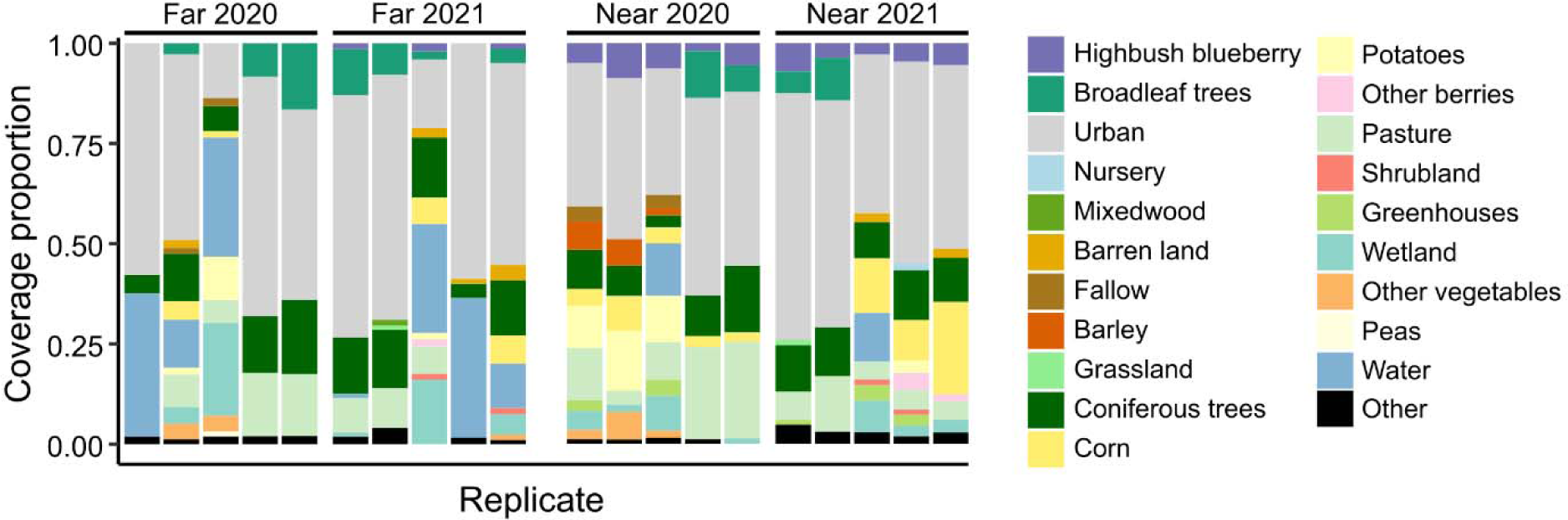
Landscape coverage. Description of land cover categories within a 1.5 km radius of t2 sites. Only categories contributing at least 1% of total coverage are shown. Data were obtained from 2020 and 2021 Annual Crop Inventory (Agriculture and Agri-Food Canada 2020, 2021).

### Multivariate community analysis

Nondimensional scaling plots of the pathogen profiles at each time point show that data points appear to cluster most strongly by year (**Figure 3**). To investigate multivariate profiles further, we first checked if profiles in replicates destined to be moved to exposure yards (n = 10 each across years, representing 80 contributing colonies) were similar at t1, before being moved to sites near to and far from highbush blueberries, and, as expected, found no significant differences (PERMANOVA, index = Jaccard; Site type: F = 0.89; df = 1, 17; p = 0.54; Year: F = 1.4; df = 1, 17; p = 0.18). Evaluating profiles at t2, t3, and t4 together, we identified a significant interactive effect of site type and time point (PERMANOVA, index = Jaccard; Interaction: F = 1.7; df = 2, 53; p = 0.021) as well as significant main effects of site type (F = 1.4; df = 1, 53; p = 0.013), time point (F = 1.6, df = 2, 53, p = 0.031), and year (F = 2.1; df = 1, 53; p = 0.021). Post hoc comparisons within each time point show that the site type effects are driven by differences at t4, the only individual time point for which pathogen profiles are significantly linked to site type (F = 2.2; df = 1, 17; p = 0.028) and year (F = 3.0; df = 1, 17; p = 0.004).

**Figure 3.**
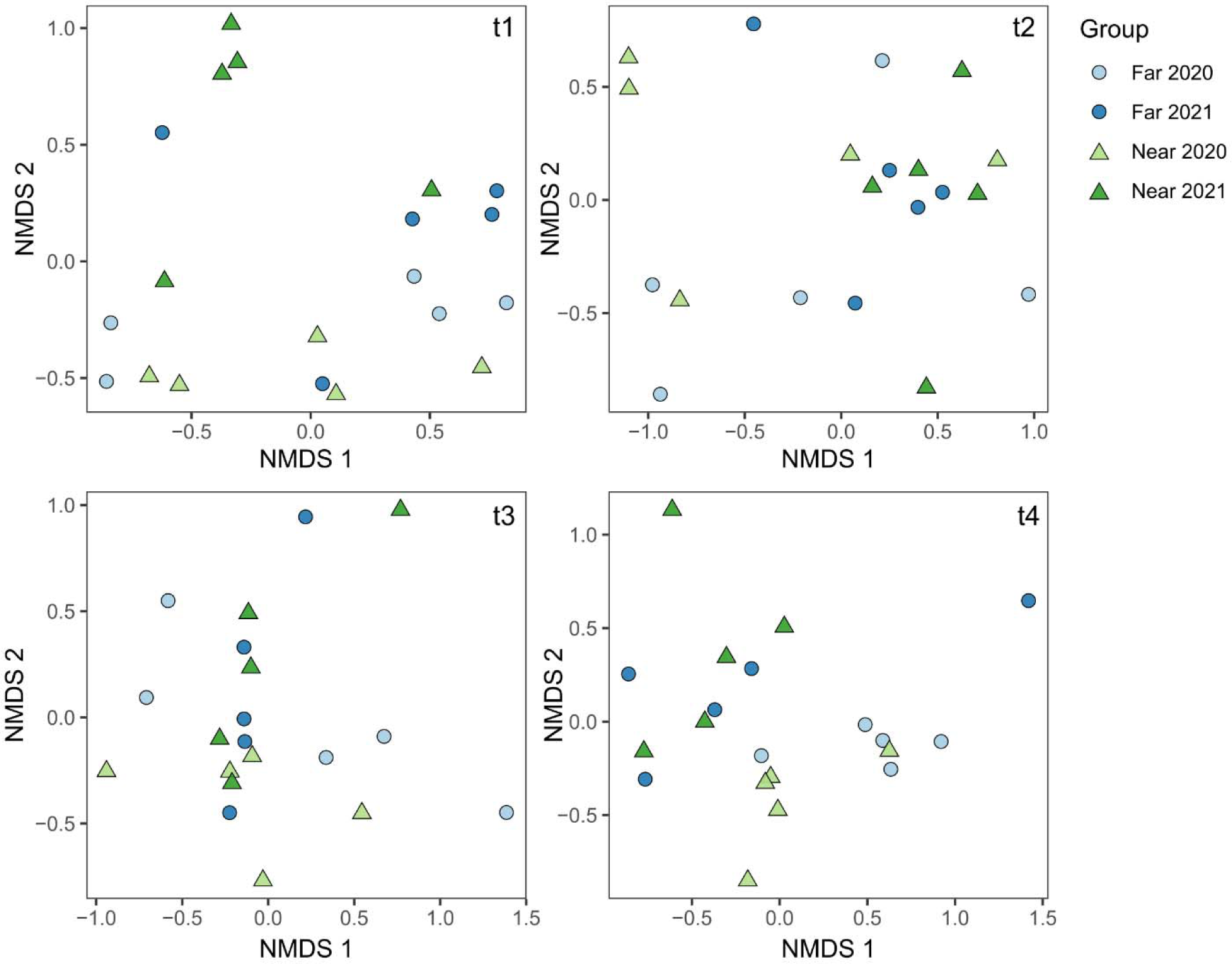
Non-dimensional scaling plots illustrating honey bee pathogen matrices. Overall differences in multivariate pathogen profiles before (t1), during (t2), after (t3), and at the end (t4) of highbush blueberry pollination were evaluated using a PERMANOVA (Jaccard index) with site type (levels: control and near) and time point (levels: t2, t3, and t4) as interactive effects, year (levels: 2020 and 2021) as a fixed factor, and replicate as a blocking factor to account for repeated measures. Data originated from N = 20 replicates (each representing a pooled sample from 4 colonies) distributed across site types and years. *Post hoc* PERMANOVA tests at each time point show that the effects are driven by t4, in particular, the separation of near and control groups from 2020.

Next, we evaluated detections of each pathogen at t4 to determine what individual profiles were driving the site type effects (logistic regression; fixed factors: site type and year) (**Figure 4**). Sacbrood virus was the pathogen most significantly linked to site type (near versus far) (χ^2^ = 8.2; df = 1; p = 0.0042; α/n = 0.0045 for Bonferroni correction), with higher frequency of detections in the samples near highbush blueberries. The next leading pathogen linked to site type was *V. apis*, but differences were not significant with Bonferroni correction (χ^2^ = 5.2; df = 1; p = 0.022; α/n = 0.0045).

**Figure 4.**
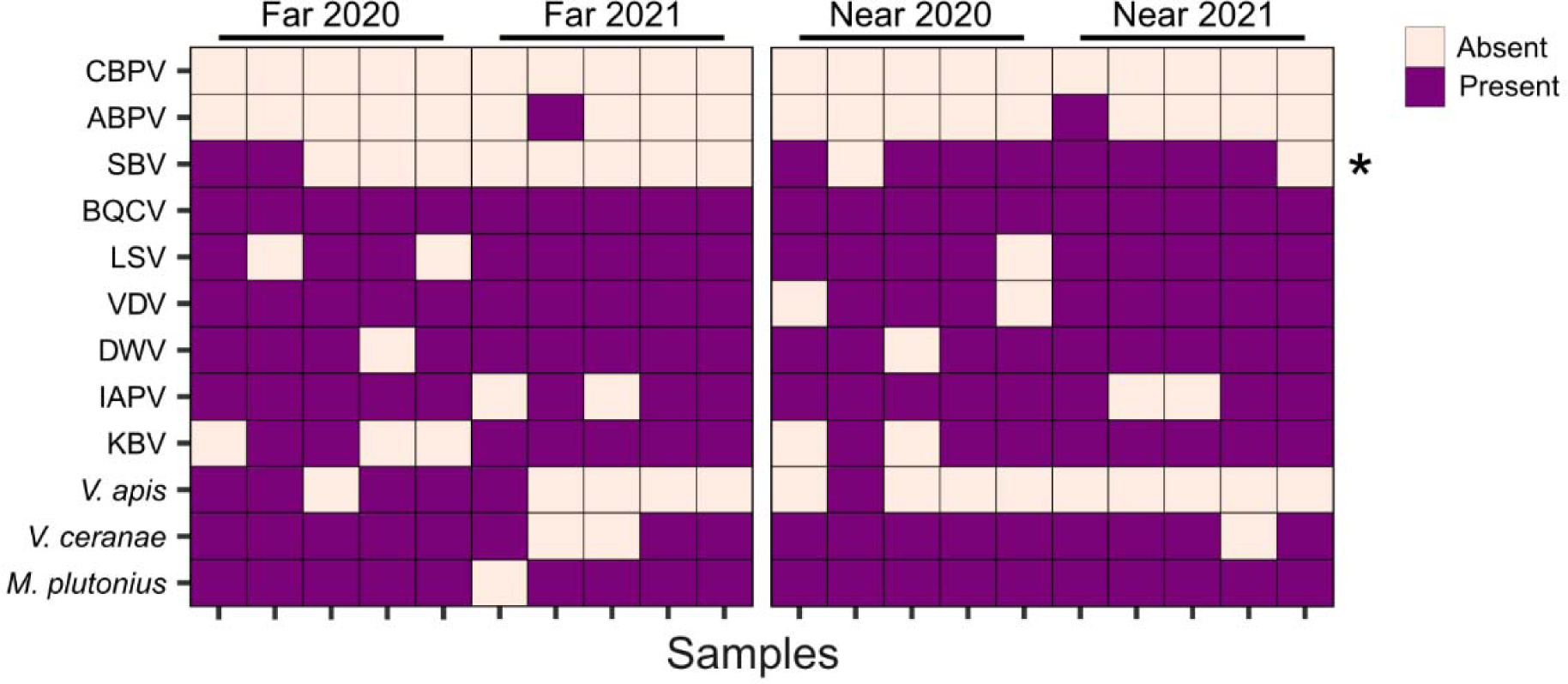
Pathogen detections at time point t4. Pooled samples (4 colonies per sample, n = 5 samples per condition, per year) were taken approximately 6 weeks after the beginning of the highbush blueberry pollination period. All pathogens were detected with qPCR or PCR. CBPV = chronic bee paralysis virus, ABPV = acute bee paralysis virus, KBV = Kashmir bee virus, IAPV = Israeli acute paralysis virus, VDV = varroa destructor virus (also known as DWV-B), SBV = sacbrood virus, LSV = Lake Sinai virus, and BQCV = black queen cell virus. Asterisks indicate significant differences determined by logistic regression with site type and year as fixed factors. BQCV was present in all samples at t4 and thus not included in post-hoc comparisons, but is still visualized here.

### Assessment of M. plutonius and SBV over time

While we did not identify a significant effect of site type at any time point except t4 in the multivariate pathogen analysis, we were still interested in patterns of *M. plutonius* and SBV detections over time, specifically. We found that *M. plutonius* detections increased over time (χ^2^ = 6.18; df = 3; p = 0.013), with no effect of site type (χ^2^ = 0.14; df = 1; p = 0.71) and, although detections in near samples did tend to increase at a faster rate than far samples, the interaction term was not significant (χ^2^ = 3.04; df = 1; p = 0.081; generalized linear model; factors: site type, time point, and their interaction; random variable: pooled sampling unit) (**Figure 5**). Using the same model structure, we found that SBV detections were not significantly linked to site type (χ^2^ = 2.98; df = 1; p = 0.084), time point (χ^2^ = 0.14; df = 1; p = 0.71) or their interaction (χ^2^ = 2.54; df = 1; p = 0.11). The significant effect of site type identified at t4 is therefore not sufficiently strong to drive an effect in this larger model.

**Figure 5.**
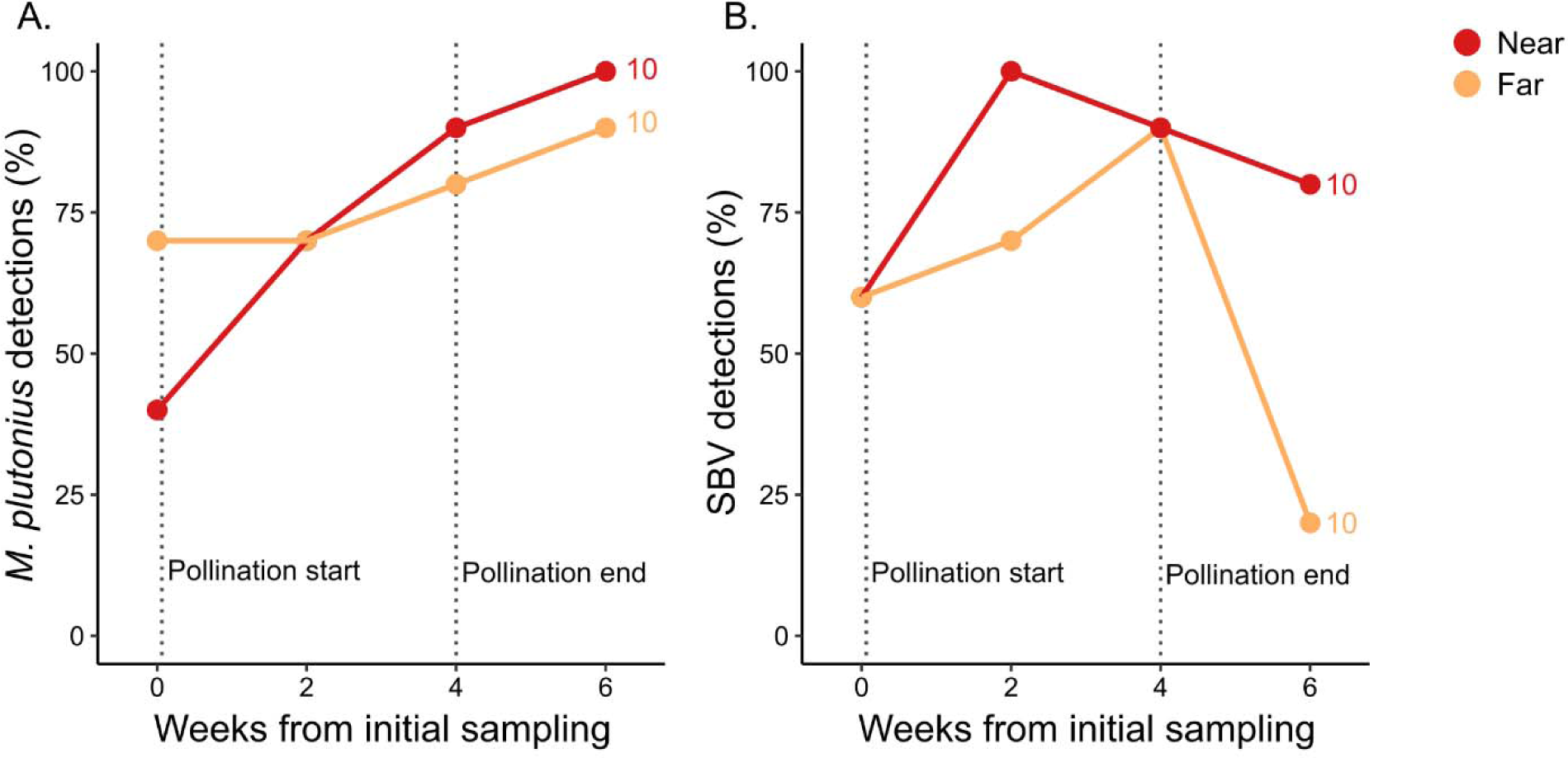
Percent prevalence of *M. plutonius* and SBV at near versus far sites over time. A) *M. plutonius* and B) SBV percent prevalence in replicates near and far (n = 10 each) from highbush blueberries. There was no effect of year for these pathogens so data from 2020 and 2021 are pooled to show overall trends. Numbers above dots indicate replicates, each representing a pooled sample from four colonies. *M. plutonius* detections are significantly linked to time point (χ^2^ = 6.18, p = 0.013) but no other significant relationships were identified (logistic regression, factors: site type, time point, and their interaction). Weeks from initial sampling are approximate and correspond to 14, 28, and 44 days in 2020 and 17, 28, and 40 days in 2021.

## Discussion

There has been ongoing concern among beekeepers in British Columbia and elsewhere in North America that engaging in highbush blueberry pollination may lead to higher prevalence of EFB disease in their colonies (Wardell 1982, Higo et al. 2019). In this study, we evaluated *M. plutonius* detections as well as nine viruses and two microsporidian parasites in honey bee colonies located far from and near to highbush blueberry fields. In a top-down approach, we analyzed all pathogen and parasite variables together (PERMANOVA, Jaccard index) and identified a significant interaction between blueberry exposure and time point.

*Post hoc* testing showed that this interaction was driven by differences at t4, and analysis of individual pathogens revealed SBV, not *M. plutonius*, to be the main driver of this pattern. Our data are consistent with Fowler *et al*. (2023), who found no relationship between EFB prevalence and blueberry pollination in Michigan. Investigating patterns of SBV over time, we found that at the end of the pollination period, both near and far groups had the same SBV detection frequency (90% of colonies tested positive), but at t4, the far group dropped to 20% while the near group remained high at 80%. This suggests that the differences observed at t4 are due to far colonies disproportionately recovering from prior SBV infections compared to near colonies. Some potential mechanisms underlying these patterns are hereupon discussed.

Like other honey bee viruses, SBV can spread between colonies as a result of robbing, drifting, and possibly through contact between bees at forage sources (Chen et al. 2006, Alger et al. 2019) – the latter two of which could be amplified to some degree when colonies are moved into pollination yards.

However, the near and far colonies in this study had the same number of SBV detections at t3, the end of the pollination period, so the difference we found at t4 is not driven by dispersal. Previous research has identified a link between fungicide exposure and increased susceptibility to some viral infections (Degrandi-Hoffman et al. 2015, O’Neal et al. 2019); for example, Degrandi-Hoffman *et al*. (2015) found that boscalid and pyraclostrobin exposure affected DWV levels more than BQCV. We did not analyze agrochemical data here, but this concept offers a potential mechanism for how some viruses may be more affected than others by crop exposure. To our knowledge, interactions between SBV infections and agrochemical exposure has not yet been investigated, but it is possible that the colonies near highbush blueberry fields were less able to clear existing SBV infections due to differences in prior agrochemical exposure.

Within colonies, SBV is transmitted horizontally (between adult workers through trophallaxis, for example), vertically (from queen to offspring via eggs), and diagonally (from nurse to larva during feeding, or larva to nurse during hygienic behavior) (Chen et al. 2006, Wei et al. 2022). As with other viruses, *V. destructor* mites act as a vector, and we were somewhat surprised to detect no relationship between mite loads and SBV detections. However, previously reported correlations are, while significant, not strong (with Pearson coefficients of 0.17 and 0.24, for example) (Borba et al. 2022). We speculate that our pooled sampling approach, smaller sample sizes, and differences in seasonal timing of samples compared to Borba et al. (2022) contribute to our observed lack of relationship between mite loads and SBV.

While our experimental design allowed us to investigate patterns of pathogen prevalence in relation to the regional timing of blueberry pollination, it does have limitations. As noted, one limitation in this study is that our statistical power was low. While the effective sample size is relatively small (n = 5 unique replicates per year, per site type), since each replicate represents a pooled sample of 4 colonies, this means that a total of 80 colonies participated in this study across years. Despite this limitation, the magnitude of the SBV effect was still large enough to detect at t4. We and others have anecdotally observed that colony health tends to decline around this time point after engaging in blueberry pollination, and we argue that SBV may be an underappreciated pathogen contributing to this observation.

A second limitation is that the high prevalence of highbush blueberry fields in the region limited our ability to place colonies in the far group such that highbush blueberry fields were completely outside of the foraging range. Average foraging distances from colonies reported in the literature is variable, ranging from 0.4-0.6 km (Schneider and McNally 1993), 0.5-1.1 km (Waddington et al. 1994), 1.1-1.4 km (Schneider and Hall 1997), 1.2 km (Schneider 1989), 2.3 km (Visscher and Seeley 1982), and 5.5 km (Beekman and Ratnieks 2000), for example. However, foraging ranges may be reduced to less than 1 km in areas of intense agricultural production, where flower resources are high (Couvillon et al. 2014, Balfour and Ratnieks 2017).

In our case, we chose the near and far field sites we did for two main reasons: 1) blueberry occurrence in the study location is so high that finding field sites outside of longer foraging radii (*e.g*., where 95% of foraging activity would take place) was impractical, and 2) the farther the distance between sites, the larger the differences in other variables would be (*e.g*., other landscape and land use parameters, microclimates, density of other beehives, *etc*.). Although some contact with blueberries may have occurred at far sites (three far study sites had highbush blueberries making up ≥1% of land cover within a 1.5 km radius), the chosen sites strike a balance between proximity to blueberries and minimizing extraneous variables. Moreover, the near sites have the key difference of being located within or immediately next to blueberry fields, as opposed to far sites for which the immediate vicinity around the colonies is blueberry-free; therefore, the near sites were still more exposed.

In this study we did not measure prevalence of EFB disease symptoms in colonies, which is distinct from detections of *M. plutonius*, as testing positive for *M. plutonius* does not necessarily mean that the colony is symptomatic (Milbrath et al. 2021). Additionally, since larval samples were not part of this experiment, we are unable to determine to what extent SBV may or may not contribute to disease manifestation or appearance. Future experiments investigating *M. plutonius* and blueberry pollination should include SBV analysis of symptomatic and asymptomatic larvae to better understand the possible relationship between these two pathogens and disease presentation, as there is some degree of overlap in their symptoms.

Like EFB, SBV symptoms are thought to occur most frequently in the spring (Bailey 1969), and like EFB and American foulbrood (AFB), dried SBV-infected larvae can also have a scale-like appearance and larvae may die after cell capping, which can lead to a similar presentation of spotty brood patterns and perforated cell caps (Grabensteiner et al. 2001, Milbrath 2021, Milbrath et al. 2021). But unlike these two bacterial diseases, SBV can replicate in adult bees and decrease their lifespan (Wang and Mofller 1970, Bailey and Fernando 1972). SBV is highly prevalent in Canada (National Bee Diagnostic Center 2017), and SBV levels in adult bees sampled in fall are associated with smaller fall and spring cluster sizes (Borba et al. 2022) as well as increased winter mortality of colonies (Desai and Currie 2016). This may in part explain the delayed appearance of site type effects in blueberry pollination units, if cascading effects of shorter-lived adults are influencing susceptibility to subsequent opportunistic pathogens.

All this is not to say that SBV detections are higher as a result of blueberry pollination, specifically. Stressors affecting disease prevalence may also originate not only from the pollinated crop but also from surrounding landscapes. Indeed, pesticide risk associated with blueberry pollination may not be driven by the crop itself, but by other crops present in the surrounding landscape (Graham et al. 2022). We speculate that there could be a broader effect of agricultural landscape exposure in general, the influence of which may or may not manifest depending on the presence or absence of additional extraneous variables.

SBV, amongst a plethora of other stressors, is generally not considered to be a major concern for honey bee health in North America. However, our data suggest that it may be an underappreciated pathogen. Relatively little research has been conducted on SBV relative to, *e.g.*, *M. plutonius*, *Paenibacillus larvae* (the agent causing American foulbrood disease), *Vairimorpha spp.,* and DWV. Given that we show significant associations between SBV detections and highbush blueberry exposure, our findings suggest that this is an agriculturally-relevant virus that deserves further attention.

## Author Contributions

Conceptualization – SEH, RWC, LJF, MMG, IMC, SF, SFP, EG, PG, AZ

Data curation – IMC, SF, PWV, MP

Formal analysis – AM

Funding acquisition – SEH, RWC, LJF, MMG, IMC, SFP, LT, EG, AZ

Investigation – PWV, JC, HH

Methodology – RWC, MMG, IMC, SFP, HH, NT, MP, AZ

Project administration – RWC, LJF, MMG, IMC, SFP, LT, AZ

Resources – JC, HH

Supervision – LJF, IMC, MP, AZ

Visualization – AM

Writing – original draft – AM

Writing – review & editing – SEH, RWC, LJF, MMG, SF, SFP, LT, EG, PG, NT, AZ

## Acknowledgements

We would like to thank Abigail Chapman, Bradford Vinson, Rhonda Thygesen, and Renee Teo for assisting with sampling honey bee colonies in BC, Bradford Vinson and Renee Teo for conducting mite washes for BC colonies. We also deeply appreciate the crop growers that cooperated with this study by providing field sites.

This work was conducted as part of the BeeCSI project, which was funded and supported by the Ontario Genomics Institute (OGI-185), Genome Canada (LSARP #16420), the Ontario Research Fund, Genome Quebec, Genome BC through the Genomic Innovation for Regenerative Agriculture, Food, and Fisheries program, and the Government of Canada through Agriculture and Agri-Food Canada (AAFC) Genomics Research and Development Initiative (GRDI) funding (AAFC J-002368).

## Data Availability

All data presented in this manuscript are available in **Supplementary Data 1 and 2.** Supplementary Data 1 includes landscape coverage data for t2 sites. **Supplementary Data 2** includes all pathogen and parasite data recorded for in near and far replicates. Primer sequences for PCR detections are included in **Supplementary Data 3**.

## Supplementary Data

All supplementary data indicated in the data availability statement have been uploaded as separate Excel files through the manuscript submission portal. If you wish to inspect the supplementary data, please refer to those, and not to subsequent pdf tables (which are automatically rendered by the submission system).

